# Software-aided workflow for predicting protease-specific cleavage sites using physicochemical properties of the natural and unnatural amino acids in peptide-based drug discovery: Peptide cleavage sites prediction workflow

**DOI:** 10.1101/340604

**Authors:** Tatiana Radchenko, Fabien Fontaine, Luca Morettoni, Ismael Zamora

**Affiliations:** Pompeu Fabra University, Barcelona Spain; Lead Molecular Design, S.L, Sant Cugat del Vallés, Spain; Molecular Discovery Ltd, London, UK

## Abstract

Peptide drugs have been used in the treatment of multiple pathologies. During peptide discovery, it is crucially important to be able to map the potential sites of cleavages of the proteases. This knowledge is used to later chemically modify the peptide drug to adapt it for the therapeutic use, making peptide stable against individual proteases or in complex medias. In some other cases it needed to make it specifically unstable for some proteases, as peptides could be used as a system to target delivery drugs on specific tissues or cells. The information about proteases, their sites of cleavages and substrates are widely spread across publications and collected in databases such as MEROPS. Therefore, it is possible to develop models to improve the understanding of the potential peptide drug proteolysis. We propose a new workflow to derive protease specificity rules and predict the potential scissile bonds in peptides for individual proteases. WebMetabase stores the information from experimental or external sources in a chemically aware database where each peptide and site of cleavage is represented as a sequence of structural blocks connected by amide bonds and characterized by its physicochemical properties described by Volsurf descriptors. Thus, this methodology could be applied in the case of non-standard amino acid. A frequency analysis can be performed in WebMetabase to discover the most frequent cleavage sites. These results were used to train several models using logistic regression, support vector machine and ensemble tree classifiers to map cleavage sites for several human proteases from four different families (serine, cysteine, aspartic and matrix metalloproteases). Finally, we compared the predictive performance of the developed models with other available public tools PROSPERous and SitePrediction.

## Introduction

Proteolytic enzymes play critical role in many processes including cell proliferation, immune response, cell death and others [20]. Protease specificity was studied in different studies [21, 36]. Protease cleavage of peptides is directed by short amino acid motifs, from two to eight amino acids around the scissile bond (site of cleavage, SoC) [17]. This specific amino acid sequence is recognized by the active site of a given protease, but the efficiency of a proteolytic activity is also related to the structural properties of the SoC. Kazanov et al. [15] studied the structural preferences of cleavage sited by mapping 200 proteolytic events to the CutDB [7]. Recently, it has been shown that the following additional factors to the peptide sequences influence on a cleavage such as unfolding events, allosteric effects, solvent accessibility, secondary structure of the sequence etc. [3]

Nowadays it is known that the new peptide-based drugs must be designed considering the localization of potential protease SoCs to make the compound less susceptible to protease reaction. Firstly, efforts are spent on evaluating a large numbers of lead compounds and their derivatives to reveal motifs recognized by proteases. Secondly, peptide sequences of potent drug candidates are typically chemically modified to eliminate these motifs or to incorporate structural features preventing proteolysis [38]. Various chemical strategies have been developed to overcome the low availability issue of peptides, including cyclization, changing the stereochemistry of an amino acid, substitution of natural amino acids by unnatural ones and others [41]. These changes are applied during the design-make-test drug discovery cycle, with hopes of improving the physicochemical and pharmacokinetics properties of the compound of interest. Therefore, it is crucial to evaluate these properties rapidly in early development and suitable in vitro techniques providing reliable predictions of in vivo performance are required.

The information about proteases and their sites of cleavage is widely spread across publications and databases such as MEROPS [26], CutDB [14] and Proteasix [7]. The MEROPS database integrates available information about proteolytic sites and, consequently, proteases from different organisms, their experimentally identified or predicted SoCs with their sequences, peptidase substrates and inhibitors. This information can be used to develop models to identify possible labile residues in the candidate peptide for individual proteases. Although useful, these databases still have several limitations. For example, none of the available resources allow to add new information in an automatic way to enrich the database information.

Different bioinformatic tools were developed to for protease-specific substrates and their cleavage sites prediction. Efficient computational tools potentially can help to reduce the number of experiments and improve peptide drug structure before synthesis. These approaches use as an input data extracted from databases described above. Several reviews were published to compare developed models and their predictive performance [2, 19, 33]. These approaches can be classified in four main groups depending on the way how models were developed. The first group contains sequence-based approaches (ExPaSy [11, 12]). The second group consists of approaches that perform prediction using position-specific scoring matrix (PSSM) for individual proteases (GraBCas [1], CasPredictor [10], PoPS [5], SitePrediction [37]). Approaches in the third group use machine learning-based predictive models trained on sets of cleavage site specific descriptors (PROSPER [30], PROSPERous [31], Pripper [23], CasCleave [34], CasCleave 2.0 [39], CASVM [40], iProt-Sub [32]). Finally, the forth group contains approaches that combine methods explained. For example, Proteasix can perform matching against the collection of the known cleavage sites from the literature and calculate probability of cleavage event appearance based on MEROPS specificity matrix.

In accordance with comparison reported in the literature machine learning methods overperform the methods from the first and the second group. In this group approaches models are trained using different features: binary features, PSSMs, molecular descriptors, structural features such as the secondary structure of the cleavage sites, the solubility of the sequence around the cleavage site and others. The cleavage site specific molecular descriptors are used to explain physical, physicochemical, pharmacophoric properties of the residues around the cleavage site. One more feature that is considered during model training is the size of local window around the cleavage site, typically from two to sixteen residues. Different machine learning algorithm such as support vector machine (SVM) learning algorithm (CASVM, Pripper), support vector regression (SVR) (CasCleave 2.0, PROSPER), logistic regression (PROSPERous), neural networks [42] and others are applied to implement protease-specific models.

This article presents a new approach which aim is to derive cleavage site appearance rules for the specific peptide family or for specific experimental condition (i.e. individual protease). The system considers the following steps: firstly, the information coming from liquid chromatography mass spectrometry (LC-MS) based experimental data or from external sources such as MEROPS database is stored in a chemically aware database (i.e. WebMetabase [6]). We represented each amino acid as a vector of physicochemical properties. We used validated Volsurf molecular descriptors [7] to describe these properties. Finally, the cleavage site-specific descriptors are calculated as a combination of the individual amino acid descriptors for the residues surrounding the cleavage site. Each stored peptide substrate and cleavage site are described as a combination of pharmacophoric and physicochemical properties of each amino acid contained in the sequence. This database can be used to perform frequency analysis (FA) to discover the most frequent SoC [7] within the experimentally derived and/or public database. FA results can be used to create a set of empirically derived rules based on molecular properties of the cleavage sites. At the next step, results of the frequency analysis for several individual proteases were used to train a Logistic Regression (LR), Support Vector Machine (SVM) and Ensemble Trees (ET) classifier prediction models. We trained models for eighteen proteases from four protease families: serine, cysteine, aspartic and matrix metalloproteases.

Nevertheless, the proposed methodology could be applied in the case of non-natural amino acid and/or cyclic peptides. Moreover, since the system used to derive the cleavage site appearance rules (frequency analysis, FA) could be linked to the software assisted metabolite structure elucidation based on MS data, the database is automatically enriched with the new experiments. Rules can then be refined to tune the system for the experimental conditions and/or peptide families of interest. This knowledge can be applied during the design-make-test drug discovery cycle.

## Materials and Methods

### In-silico MEROPS dataset

In this study we exported peptide cleavage data from the MEROPS database (version 11 1/09/2017) [26] for all the available proteases in MySQL format using our own script that converts each organism/protease/peptide information in a single xml file inside a folder data structure. This data consists of substrate sequences, the cleaving protease, and the cleavage sites. MEROPS database version 11 contained information about 75959 substrates. Each peptide sequence was converted to a chemical structure and saved in MDL mol format. [9] For our analysis, we focused on peptides having less than 200 amino acids. This amount of the amino acids was selected since we were concentrated on peptide substrates more than proteins. Several peptides were excluded from analysis because it was not possible to restore their chemical structure. The reasons could be, for example, the lack of an appropriate defined structure for some encoded abbreviations of amino acids in a sequence.

Two different strategies were followed depending if the source organism for the protease was indicated in the MEROPS database or it was not. The summary information extracted from the MEROPS database includes: substrate peptide sequence, parent sequence peptide range, substrate Uniprot number, cleavage site position in substrate, metabolite peptide sequence, protease name, protease source organism and references list related to the protease. All the information was collected and written in a xml file. The xml has four blocks of information:

1. Properties: The Matrix (name of the protease), the MEROPS Peptidase Code, the MEROPS Peptide Uniprot_acc, the MEROPS Peptide Range, the MEROPS Organism and the MEROPS Peptidase Reference.
2. Parent: the parent structure in sdf format, the InchiKey, the molecular formula and the m/z.
3. Metabolized Parent: as the previous one without the InchiKey.
4. Metabolites: There is a new block for each metabolite. It contains the metabolite name and the structure. Each structure has the structure ID inside the xml file, the sdf (V3000 or V2000 format), the formula, the m/z, the formula difference compared to the parent structure, the metabolic mechanism, the m/z difference compared to the parent, the collection of atoms in the parent that suffer the metabolic transformation and the list that contains the isomorphism between the number of the atom in the parent and the number of atom in the metabolite.

After the xml files were imported into WebMetabase the same steps to annotate the information for each file that were described elsewhere for the case of experimentally derived approach were followed [25]. Enabling in this way the combination of the external and internal sources of information.

## WebMetabase

All files extracted from MEROPS database were uploaded into the web application “WebMetabase 3.2.12” (Molecular Discovery Ltd, Middlesex, UK). WebMetabase imports the data from external sources in a predefined xml format where the structure of the peptides introduced in MDL mol format (V3000 or V2000). This conversion to a structure format is necessary to feed the chemically aware system. A complete list of uploaded substrates for each protease grouped by organism can be found at http://webmetabase.com:8182/WebMetabaseBioinformatics/. To access this database following credentials should be used: login “User”, password “WebMetabase”.

### Peptide database and search algorithm based on similarity

Once the experimental results and/or data from external database were interpreted and approved in WebMetabase, the parent and metabolite structures were stored in the database. Each peptide structure was annotated by the Structural Blocks (SBs) defined as the structural fragment between amide bonds and their connectivity. SBs in the substrate sequence were numbered as …-P4-P3-P2-P1-/-P1’-P2’-P3’-P4’-…, with the cleaved bond located between the P1 and P1’ sites. [28, 29]

In this article we described a new approach implemented in WebMetabase to store each peptide described as a combination of physicochemical properties of each SB contained in the sequence. These physicochemical properties of the SBs were represented through molecular descriptors calculated by the cheminformatics library Volsurf. [7] Therefore, each SoC P1-P1’ was also characterized as a combination of physicochemical properties of the two SBs. Moreover, similarity matrix for Volsurf descriptors was calculated for all SB stored in the database.

The annotation of the peptide information in this manner enables doing an exact substructure and similarity-based substructure search inside the database without being limited to any type of peptides and/or amino acid (cyclic/linear, natural/synthetic). This methodology can be used to perform a structural search on the exact sequences identifying the experiments where a certain bond of interest participated in a metabolic reaction. Also, the described algorithm implemented in WebMetabase includes a system to perform searches based on similarity of the molecular descriptors. As an output, searches give a list of experiments that fulfill the search criteria. [25]

### Volsurf descriptors

We used the cheminformatics toolkit Volsurf [7, 8] to calculate the physicochemical properties for the individual SBs. Peptide descriptors are then built from the descriptors of the SB in the peptide. Volsurf+ creates 128 molecular descriptors from 3D Molecular Interaction Fields (MIFs) produced by our software GRID [4, 13]. The Volsurf descriptors summarize the MIF information in a few variables easy to understand and to interpret. Basically, it quantitatively characterizes size, shape, polarity, hydrophobicity and the balance between them.

### Frequency analysis

The algorithm can perform frequency analysis (FA) based on exact match of the sequence of interest (simple frequency analysis, sFA) and/or on the similarity based on the molecular descriptors (similarity frequency analysis, SFA). A simple peptide frequency analysis of the cleaved amide bonds can be performed for the entire database or for the selected set of the approved experiments in WebMetabase. The algorithm collects information about all cleaved peptide bonds that were involved in the metabolic reactions of interest. Herein site of cleavage (SoC) considers the two SBs involved in the amide cleavage containing the C-terminal and N-terminal of the SBs. We define a potential SoC (pSoC) as the pair of structural blocks (referred as P1 and P1’) that may and may not be involved in the catalysis. The frequency analysis refers to the number of times that a pSoC is observed in the parent structure and how many times this is an actual SoC (aSoC).

The output of the frequency analysis is done by protease and provides two tables. The first table contains the counts of a pSoC found and the counts that it was an aSoC. The second table contains the counts a pSoC found and if it was cleaved once, twice or more than twice. A frequency analysis of the aSoCs depending on the protease can be done to create a set of empirically derived rules that later can be used to predict the metabolic liability of different amide bonds in a new nontested peptide. WebMetabase automatically generates an excel file in temporary folder to store FA results. Filename format starts with string “soc” and after some random number, e.g. soc7511073558888689580.xls. This file can be used for the following analysis if needed. Moreover, frequency analysis results can be persisted in the database for the further usage.

## Model implementation

The information registered in the database and FA results based on MEROPS dataset for eighteen proteases were selected to make some examples of the methodology. These proteases from four different families (serine, cysteine, aspartic, matrix metalloproteases) from Homo sapiens were chosen to develop classifier models. They included granzyme A, granzyme B, granzyme B (rodent-type), granzyme M, thrombin, trypsin1, caspase-1, caspase-2, caspase-3, caspase-6, caspase-7, cathepsin E, cathepsin D, cathepsin L, MMP2, MMP3, MMP8, MMP9. Trypsin1, MMP2, caspase-6 and cathepsin L were selected because the number of the identified aSoCs was the highest for proteases imported in WebMetabase. Others were selected as a representative of the respective protease family. For each selected protease we split all extracted substrates in train and external dataset (ED). Train dataset contained experiments with 80% of extracted cleavages events and was used to develop models and to perform the cross-validation. ED was used only as a final assessment of model predictive performance quality for each protease and to perform comparison with publicly available PROSPERous and SitePrediction approaches. In accordance with the literature it was revealed that increasing of the investigated window’s size around cleavage site could improve the performance of the model [31]. Due to this fact, all classifier models were built using two sequence windows, consisting of the P1-P1’ (SoC2) and P4-P4’(SoC8). SoC2 considers the two SBs P1 and P1’ involved in the amide cleavage containing the C-terminal and N-terminal of the SBs. SoC8 considers the eight SBs on positions P4-P4’. S1 Table summarizes the information regarding selected proteases, protease family, number of selected substrate peptides, number of cleavage events exported from MEROPS, number of unique aSoCs from two amino acids (P1-P1’), number of unique aSoCs from eight amino acids (P4-P4’) and number of substrates excluded from the exported dataset to perform external validation (EV) of the models.

To derive a training and external dataset for each of the selected proteases, we cut out all possible 2 SBs and 8 SBs sequences from substrate peptides. Thus, a substrate with 100 amino acids would lead to 93 and 99 sequences of length 8 and 2, respectively. Each sequence than was characterized as a combination of physicochemical properties described by Volsurf descriptors of the SBs in the sequence. These sequences were then classified as either cleaved (aSoC2 and aSoC8) or non-cleaved sequences (pSoC2 and pSoC8). Cleaved sequence for aSoC2 was the one involved in metabolic reaction. In cleaved sequence for aSoC8 the proteolytic event happened between fourth and fifth residue. Substrate cleavage site prediction was formulated as a binary classification problem for two classes cleaved SoCs so-called positive examples or non-cleaved SoCs so-called negative examples. We calculated number of times each pattern was cleaved and number of times it was met. For the SoC2 it was done during sFA in WebMetabase, for the SoC8 additional script was applied. Since the size of positive dataset was significantly smaller than the number of non-cleaved motifs, we split all negative dataset in groups with number of objects equal to the number of cleaved patterns, than combined positive dataset with each of the received negative subsets, therefore obtained balanced datasets to derive models. In details, we decomposed the non-cleaved objects dataset into X partitions, where X depended on the number of cleaved sequences and combined all the metabolized SoCs with each partition of pSoCs to be an individual subset. Thus, we generated multiple training subsets with better class distribution and each noncleaved object occurred in at least one training subset (Fig 1). Moreover, we performed directed oversampling for aSoCs and pSoCs to give a weight to the motifs that appeared in the dataset and were metabolized more frequently [24, 35]. We gave them an over representation in the dataset for certain number of times. This number was calculated in accordance with the following formula (2). S2 Table summarizes the information regarding number of subsets for each protease.

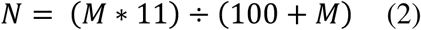

**Fig 1.**
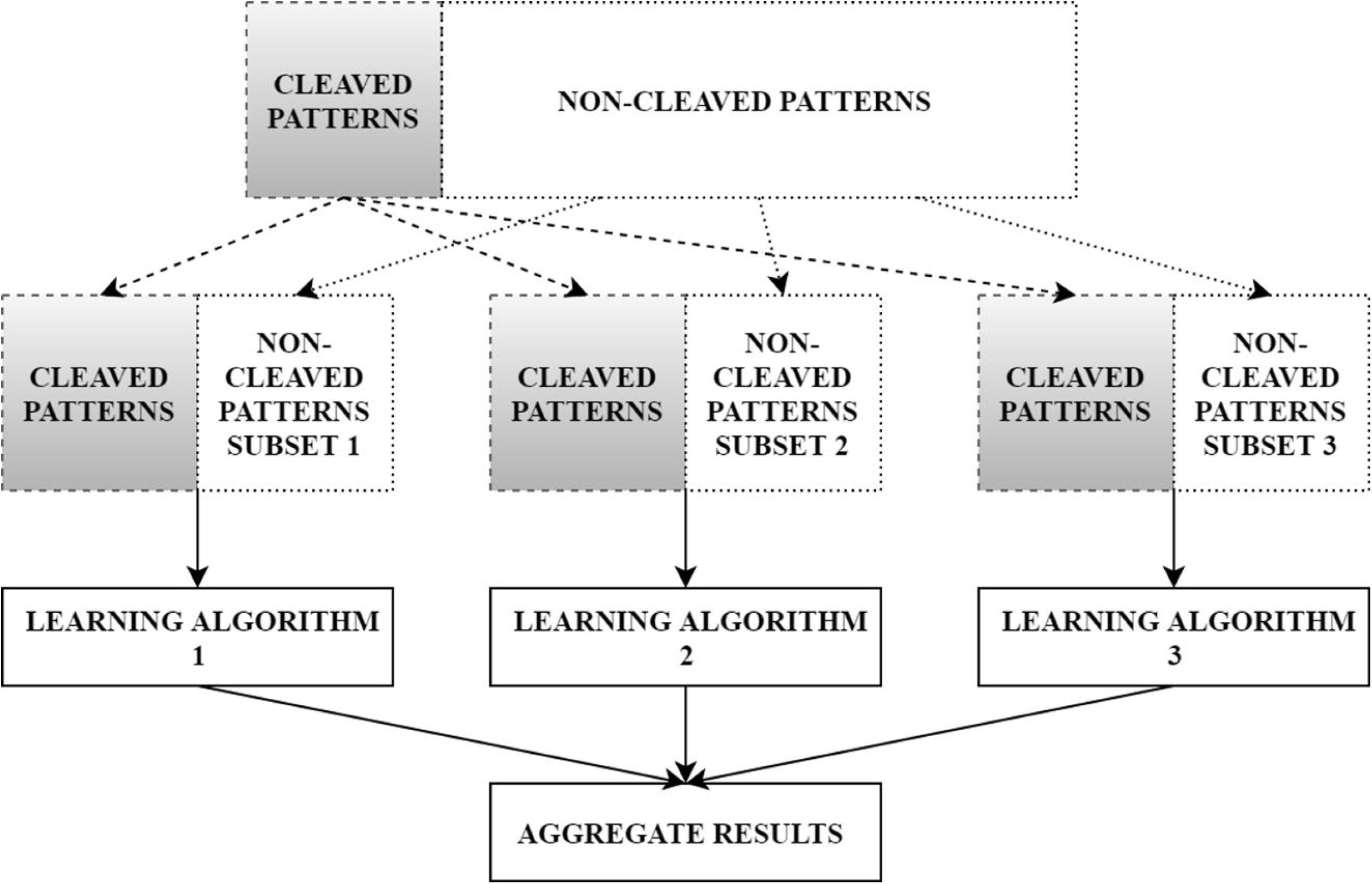
Architecture of the classifier ensembles.

N is the number of times sequence pattern will be repeat in a final dataset. M is the number of times the sequence pattern was met in the initial exported dataset. Train dataset and external dataset for each of the selected proteases used for model training for local window of two residues and local window of eight residues are provided in the Supporting information S1 File.

The next step, for each protease we applied multiple learning algorithms to each training subset of sequences described by Volsurf descriptors to train prediction models independently. In this study, we used Logistic regression (LR) [22, 43], Support Vector Machine Classifier (SVC) [2, 44] and Random Forest Classifier (RFC) [22, 45] and Gradient Boost Classifier (GBC) [22, 46] to build the models to estimate the cleavage site probability of substrates for selected proteases. For SVC we selected radial basis kernel function (RBF) [22, 47] and applied Grid Search CV [22, 48] to search best parameters of the classifier. For all other classifiers default parameters were used.

While models were generated, we performed 5-fold cross-validation (5CV). The predictive performance of each model was evaluated using the accuracy, sensitivity, specificity, area under the curve (AUC) for the receiver-operator characteristic curve (ROC), area under the curve for the precision-recall curve (PRC), Mathews correlation coefficient (MCC) using these measures based on 5-fold cross-validation. Finally, we combined all models measures for each learning algorithm by calculating the mean for each measure.

To evaluate the model’s performance in a real scenario and to inspect the potential overfitting effect on these algorithms, we also performed prediction on EDs, peptides that had not been included in the model building. Prediction was performed for each peptide from the external dataset independently by each model. To complete this goal each peptide from external dataset also was split in motifs and each motif was characterized as a combination of physiochemical properties of its SBs. To combine the results from all models for each learning method we calculated a mean of predicted probabilities for each bond in the peptide, using the predicting bond breaking probability we evaluated ranking performance for each classifier. The procedures are described in detail below.

Finally, we performed independent tests and compared the predictive performance by ranking on ED between selected LR and RFC model classifiers and the other public available tools such as SitePrediction and PROSPERous.

## Evaluation of model performance

### Performance evaluation comparison between different classifiers

We performed comprehensively evaluation and comparison of the predictive performance between different classifiers (LR, SVC, RFC and GBC) on the external dataset (ED) in each protease family combining the prediction for all proteases inside the family. Based on the intermediate results presented below we decided to proceed with classifiers using local window P4-P4’. To complete this task, we cut out each external peptide in subsequences of length 8, calculated Volsurf descriptors for each of the SB in this peptide and characterized each sequences of 8 SBs as a combination of physicochemical properties of SBs contained in the subsequence. We used these combinations of Volsurf descriptors to perform prediction on each of them. Since the final goal of the model was to select which of the bonds was more likely to be broken in each sequence, an analysis of the rank (sorted by the predicted SoC) was performed. Therefore, when the probability for each bond was computed by the model, we sorted each sequence by their value and registered a rank of each predicted cleavage site. Since peptides differ in a size ranking was normalized by the amount of possible cleavage sites. A recovery analysis of the rank value for the known SoC was done and finally we accumulated all ranking positions for each external peptide and each protease inside the family. Moreover, we calculated the best and the random ranking. The random ranking was done by assigning a random number for the probability to be broken for each SoC and continuing in the same way as in the model case. In the case of the best ranking positions, the known SoCs in a peptide were assigned to 1,2,3, etc. rank for each of the known SoCs. As a summary of the results, we plotted the cumulative ranking score for the recovered known SoCs for each family including the best and the random cumulative ranking. Python algorithm used to perform predictive models training and to perform prediction on external dataset for each model for each of the selected protease with ReadMe.txt was provided in the Supporting materials (S2 and S3 Files).

### Performance evaluation comparison with other prediction tools (SitePrediction and PROSPERous)

We performed comprehensively evaluation and comparison of the predictive performance between selected LR and RFC model classifiers and other public available tools such SitePrediction and PROSPERous. Comparison with the external methodologies was performed only for the proteases in each family where models were available. Models for trypsin1, granzyme M and caspase-2 were not available in both methods, also for granzyme A, granzyme B, granzyme B (rodent-type) and thrombin were not available in SitePrediction approach. To complete this task, we used as an input for SitePrediction and PROSPERous FASTA format sequences of the peptides in the external dataset and exported prediction results. In SitePrediction we selected models trained for each protease for Homo sapiens. In PROSPERous we selected logistic regression models and local window P4-P4’. After each potential cleavage site was sorted in accordance with the predicted scores from each method. For the SitePrediction case where a score was not predicted it was assigned to 0. After we followed the same rank strategy explained for the models developed in this article, registering the rank of each predicted cleavage site and normalized it by the amount of possible cleavage sites for each peptide. Finally, we plotted the cumulative ranking score for each family for our models and for each method including the best and the random cumulative ranking.

## Results and Discussion

In this section, we present the results of the application of our approach in the analysis of the MEROPS. All files extracted from MEROPS database were uploaded into the web application “WebMetabase 3.2.12” (Molecular Discovery Ltd, Middlesex, UK). WebMetabase imports the data from external sources in a predefined xml format. Metabolites structures extracted from MEROPS were directly uploaded into WebMetabase and automatically approved. After approval parent peptide and metabolites were persisted into the database and annotated as described before. Finally, the methodology was applied to perform a sFA on the stored information for selected proteases: granzyme A, granzyme B, granzyme B (rodent-type), granzyme M, thrombin, trypsin1, caspase-1, caspase-2, caspase-3, caspase-6, caspase-7, cathepsin E, cathepsin D, cathepsin L, MMP2, MMP3, MMP8, MMP9. The sequence pattern around the potential cleavage site was represented as a combination of Volsurf descriptors. Moreover, an additional script was applied on the database to perform sFA for the sites of cleavage with the local window P4-P4’. The results from both FA were used to build predictive models for the selected proteases enabling the cleavage site prioritization based on the physicochemical properties for the cleavage sites. The MEROPS/WebMetabase workflow used is shown in Fig 2.

**Fig 2.**
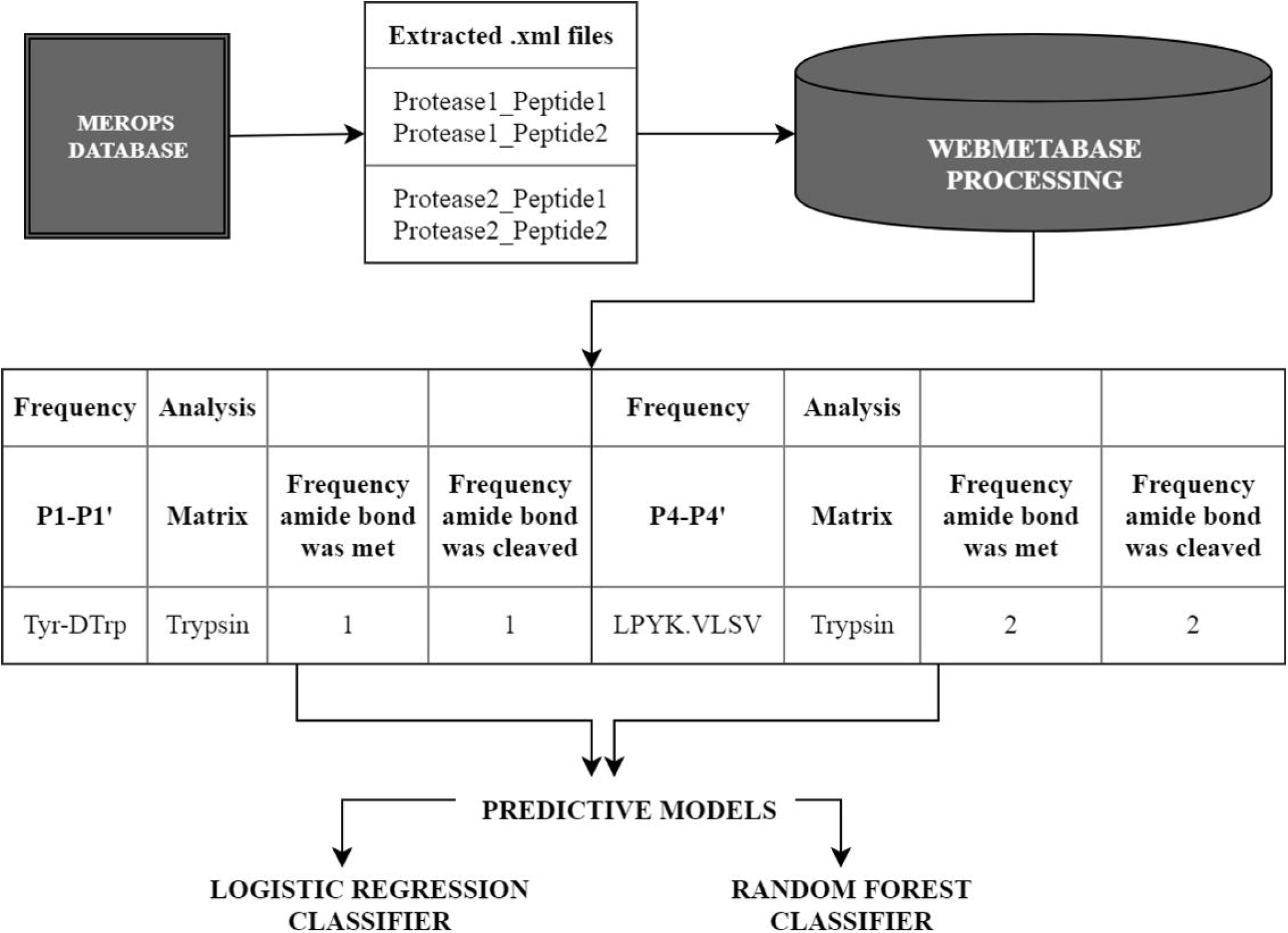
MEROPS/WebMetabase workflow from experimental data to searchable information manageable by *in silico* analysis tools.

In total we extracted 18760 xml files and they contained information for 18760 substrate peptides and 21804 metabolites. For each selected protease we split all extracted experiments in train and external dataset. Train dataset contained experiments with 80% randomly selected of the extracted identified cleavages events and it was used to develop models and perform crossvalidation. External dataset was used to evaluate the model performance and compare it with other methods. In both datasets each SoC was represented as a combination of the Volsurf descriptors of the SBs in the sequence. We performed a simple frequency analysis of the metabolized chemical moieties for each protease in the train and external datasets. The summary on the amount of identified individual metabolized bonds by frequency analysis is presented in S3 Table. Full information of identified cleavage sites frequency can be found at http://webmetabase.com:8182/WebMetabaseBioinformatics.

Using the prepared substrate training datasets, we analyzed the statistical distributions of cleavage sites for the eighteen proteases analyzed (S1 Table). The amino acid occurrences in the P4-P4’ positions for the SoCs of all selected proteases were calculated to generate heat map. The heat maps for the selected caspases and trypsin1 are presented as an example in Fig 3, for all other proteases they are provided in S1-3 Figs. These heat maps help to identify conserved and frequently occurring amino acids at scissile bonds (Fig 3, S1-3 Figs). For example, as shown in Fig 3, one of the main characteristics of caspase-3 specificity is that this protease preferentially cleaves after the aspartic acid (Asp, D) at P1 and P4 position. We identified Asp in position P1 in 100% of analyzed SoCs and in position P4 in 57.1%. Also, we noted that glutamic acid (Glu, E) presented in 28.6% at P3 and valine (Val, V) presented at position P2 in 30.95%. Also, glycine (Gly, G) at P1’ in 30.95% and alanine (Ala, A) in 23.8% or leucine (Leu, L) in 21.4% at position P2’. Therefore, a good substrate for caspase-3 should contain sequence like D-E-V-D+G-A or D-E-V-D+G-L. These results agree with literature where it was described caspase-3 preferentially cleaved at D-E-V-D+X-X-X [24, 35], Comparison of different caspases reveals differences between preferential cleavage motifs, for example, phenylalanine at P2’ for caspase-1, serine at P1’ for caspase-2 and caspase-6, threonine at P2 and leucine at P4’ for caspase-7. These results also agreed with specificity matrix presented in MEROPS DE-E-V-D+GS---. This indicates that, although we reduced our dataset by selecting only peptides of small size limited to 200 of amino acids, we still got representative dataset of substrates for each protease.

**Figure. 3.**
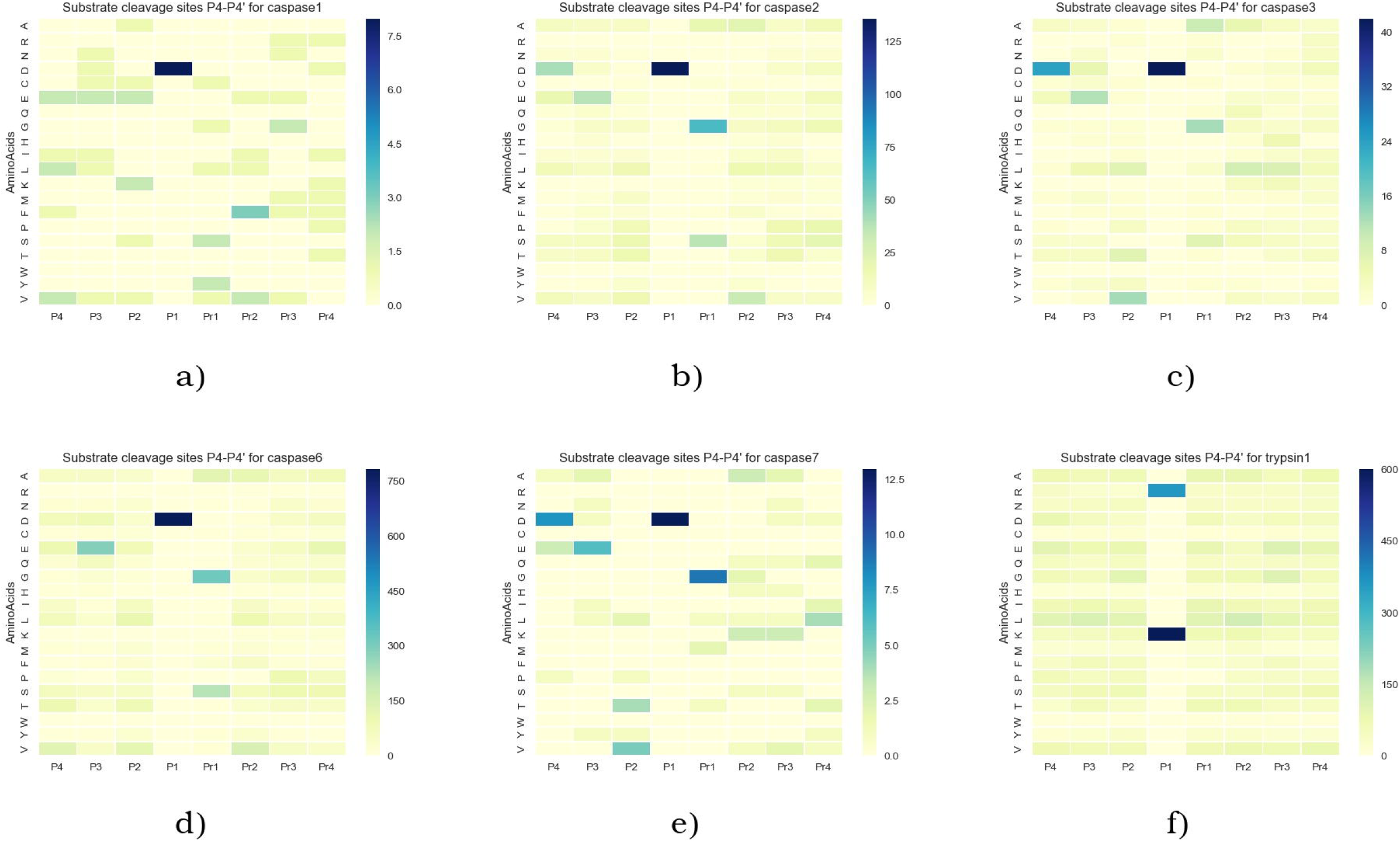
The cumulative amino acid occurrences in P4–P4’ two-dimensional heat map for caspases: a) caspase-1; b) caspase-2; c) caspase-3; d) caspase-6; e) caspase-7; f) trypsin1. The scissile peptide bond was between sites P1 and P1’.

In addition, for trypsin1 we identified arginine (Arg, R) in position P1 in 36.8% and lysine (Lys, K) in 63.2%. These results agreed with literature where it was described trypsin preferentially cleaved at Arg and Lys in position P1 [16] and with MEROPS specificity matrix.

To evaluate the performance of the selected classifier models for cleavage site prediction of multiple proteases, we carried out a 5-fold cross validation (5CV) test on each of selected proteases under investigation in this study. Moreover, the predictive performance on several external datasets was explored. Firstly, we trained and compared the predictive performance of models with two learning methods LR and SVC using two sizes of local windows P1-P1’ and P4-P4’ For evaluation of predictive performance during 5CV test the following criteria were calculated: accuracy, ROC AUC, sensitivity and specificity. For evaluation of the predictive performance on external dataset we calculated the accuracy, sensitivity and specificity. S4 and S5 Table summarizes results for the predictive performance evaluation for LR and SVC classifiers trained using local windows P1-P1’ and P4-P4’ for caspases all evaluated parameters for 5-fold CV and for external dataset, respectively. S6 and S7 Table summarizes results for the predictive performance evaluation for LR and SVC classifiers validated on external dataset for all selected proteases and all evaluated parameters trained using local windows P1-P1’ and P4-P4’, respectively.

For caspases-1, 3 and 7 all performance measures decreased for both classifiers when window was widened, on the other hand for caspases-2 and 6 they increased for LR and decreased for SVC. For example, for caspase-1 and 2 comparing LR classifier performance based on AUC ROC for the 5CV results for window P1-P1’ was 0.77 and 0.87 and for P4-P4’ was 0.73 and 0.93, respectively. Comparing results for LR and SVC specificity was higher for SVC classifier in both P1-P1’ and P4-P4’. Meanwhile, when we compared predictive performance on external dataset of the models trained using two local windows AUC ROC for P4-P4’ were higher for caspases-2, 3, 6, 7 and cathepsin L in both classifiers. Also, AUC PRC and MCC were higher for P4-P4’ were higher for caspases-2, 6 and cathepsin L in both classifiers For caspase-1, −3 and −7 AUC ROC, AUC PRC and MCC were lower that also could be related with small amount of training data. It can be explained by the fact that models trained for window P4-P4’ are more complex and involve higher number of variables to be defined during the training process. We suppose that because the amount of data is not sufficient to properly train the model for window P4-P4’. For caspases-1, 3, and-7. This theory is also confirmed by the fact that all parameters increased when P4-P4’ dataset was used to train both learning methods for cathepsin L. Based on this analysis results we decided to continue using local window sequence P4-P4’.

In addition, we compared the difference of amount of recovered known cleavage sites for all selected proteases for the P1-P1’ and P4-P4’ LR models for the 10 top positions. To complete this task, we calculated a difference and normalized it by total amount of known cleavage sites. In Fig. 4 amount of recovered known cleavage sites for 10 top positions is shown for LR models trained on P4-P4’ local window.

**Figure 4.**
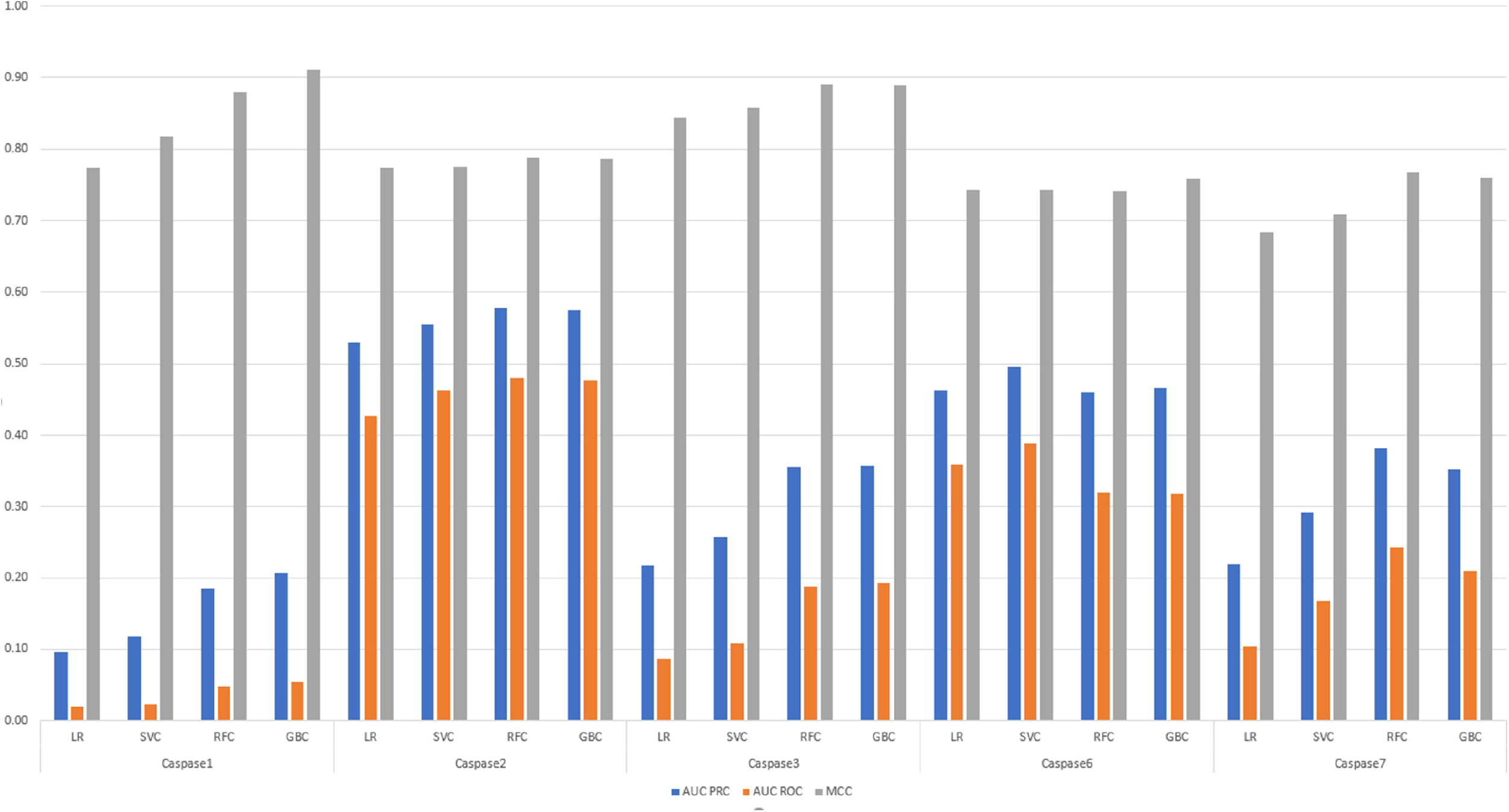
The amount of recovered SoCs for LR models trained on P4-P4’ local window for 10 top positions.

Different learning approaches methods were used to train the models in other tools [30, 31, 39, 42]. For example, in the PROSPERous approach the best performing models are based on logistic regression. [31] At the next step, we trained and compared predictive performance of models with the following learning methods: LR, SVC, RFC and GBC. Fig 5 summarizes information regarding MCC, AUC PRC and AUC ROC evaluated on external dataset for all investigated classifiers comparing predictive performance of these models for caspases using window around cleavage site P4-P4’. Both ensemble tree classifiers outperformed logistic regression and SVC for all caspases by comparing all three metrics. All evaluated metrics (accuracy, AUC PRC, AUC ROC, MCC, sensitivity and specificity) for all investigated classifiers comparing predictive performance of these models for all selected proteases using window P4-P4’ are shown in S8 Table. In addition, predictive performance of RFC and GBC models was equal for caspases-2 and 6, meanwhile RFC demonstrated higher accuracy for caspase-7 and GBC overperformed RFC model for caspase-1 that also can be related with the size of the training dataset.

**Figure 5.**
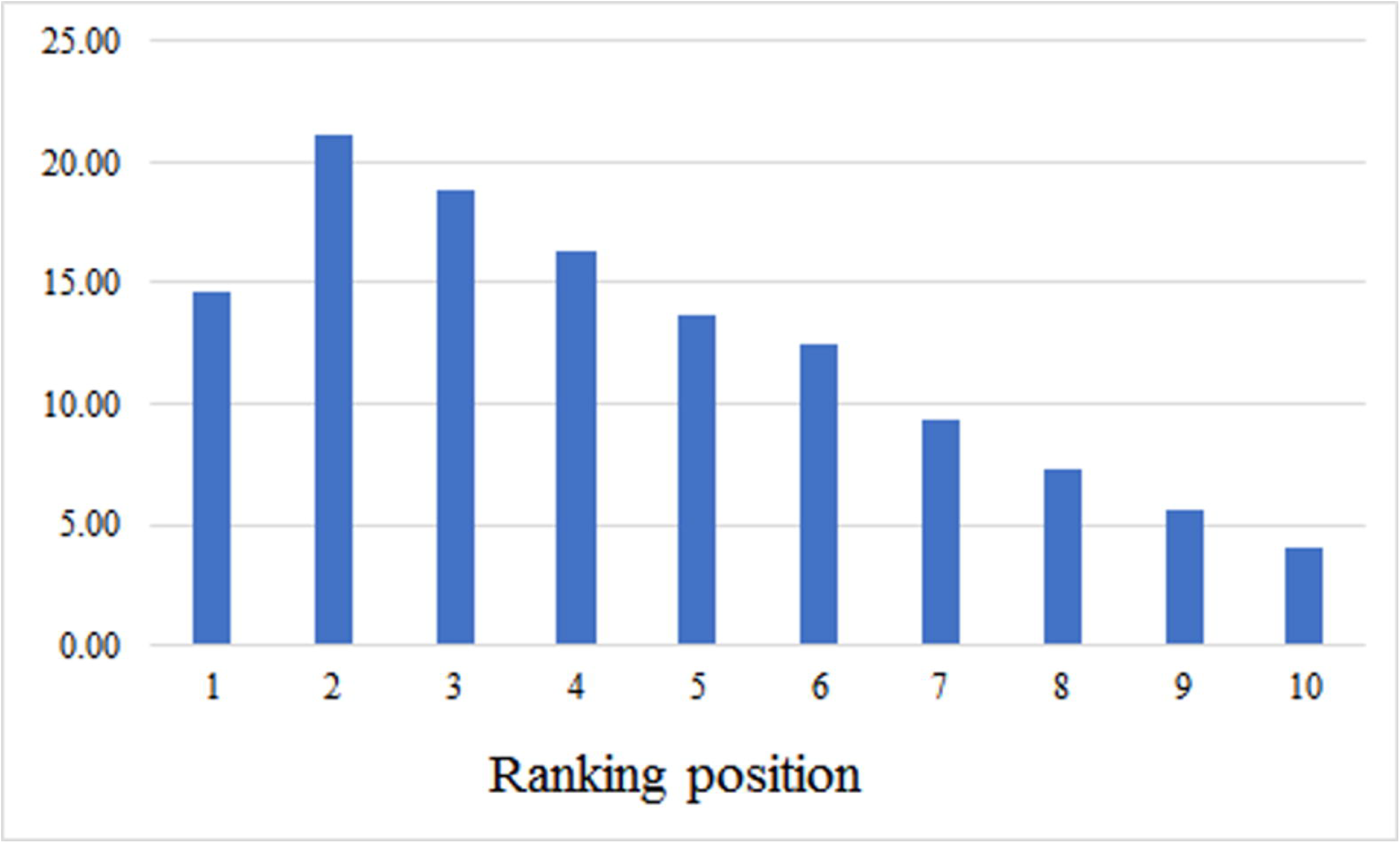
The predictive performance results comparison for MCC, AUC PRC and AUC ROC for all investigated classifiers for caspases using local window P4-P4’.

Lastly, we evaluated predictive performance of LR, SVC, GBC and RFC models on external dataset to understand if trained models were able to predict known cleavage sites with the top-ranking positions for the protease of interest. To complete this task, we explored the ranking positions of the probable cleavage sites sorted by predicted probability for each learning approach. Since peptides differ in a size, ranking was normalized by the amount of possible cleavage sites. Therefore, we completed a recovery analysis of the rank value for the known SoCs and calculated the sum of all ranking positions for each external peptide for each protease inside the family. Moreover, we calculated the best and random ranking. The cumulative percentage ranking score for the recovered sites of cleavage for all proteases is demonstrated in Fig 6.

**Figure 6.**
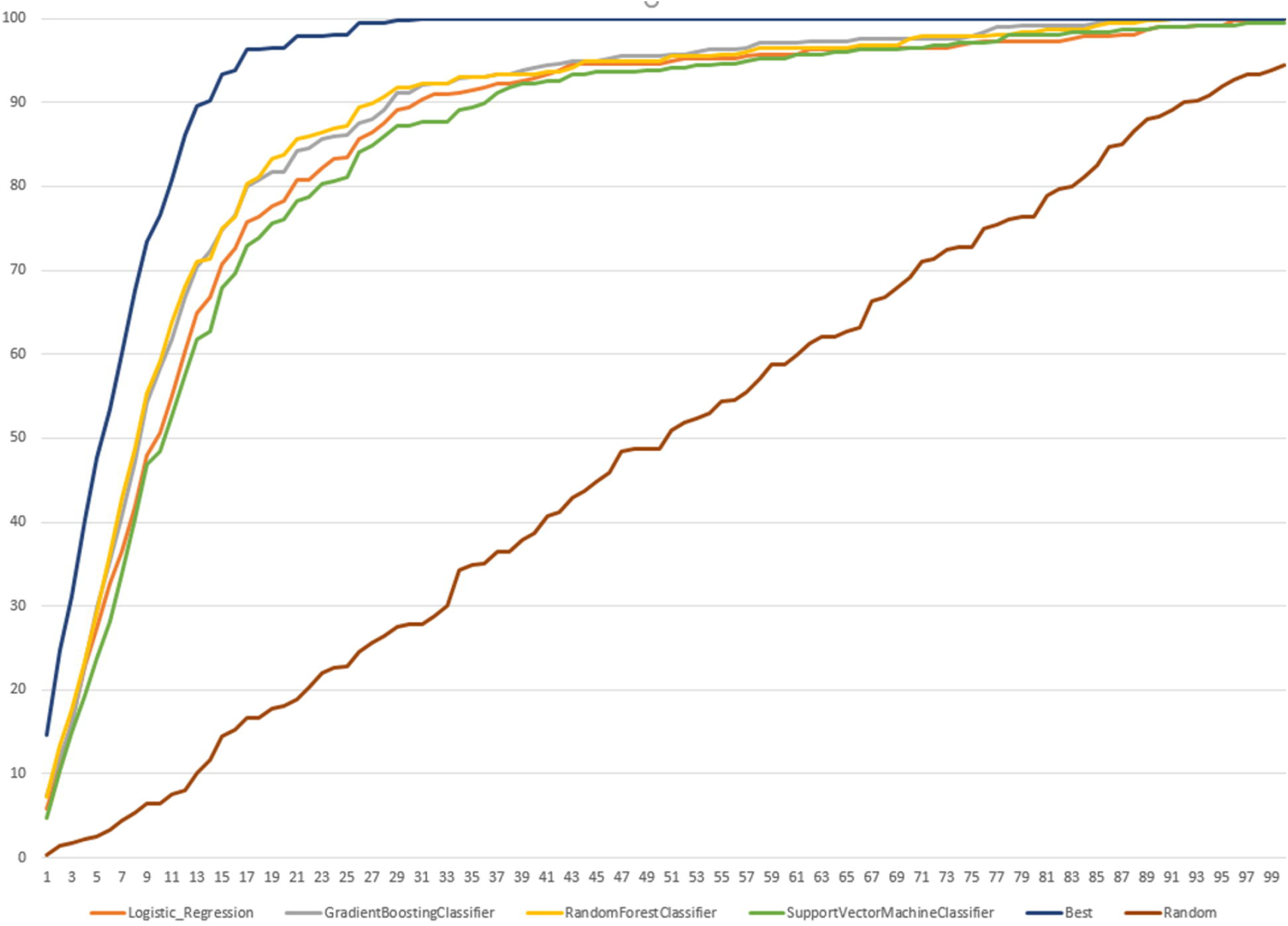
The cumulative ranking score in percentage for the recovered known sites of cleavage for all selected proteases in range of ranking positions from 0 to 100%.

We noted that the best ranking recovered 100% of known SoC reaching 20% of the normalized ranking positions. Fig 7 summarizes percentage recovered by each learning algorithm at 20% of the normalized ranking positions.

**Figure 7.**
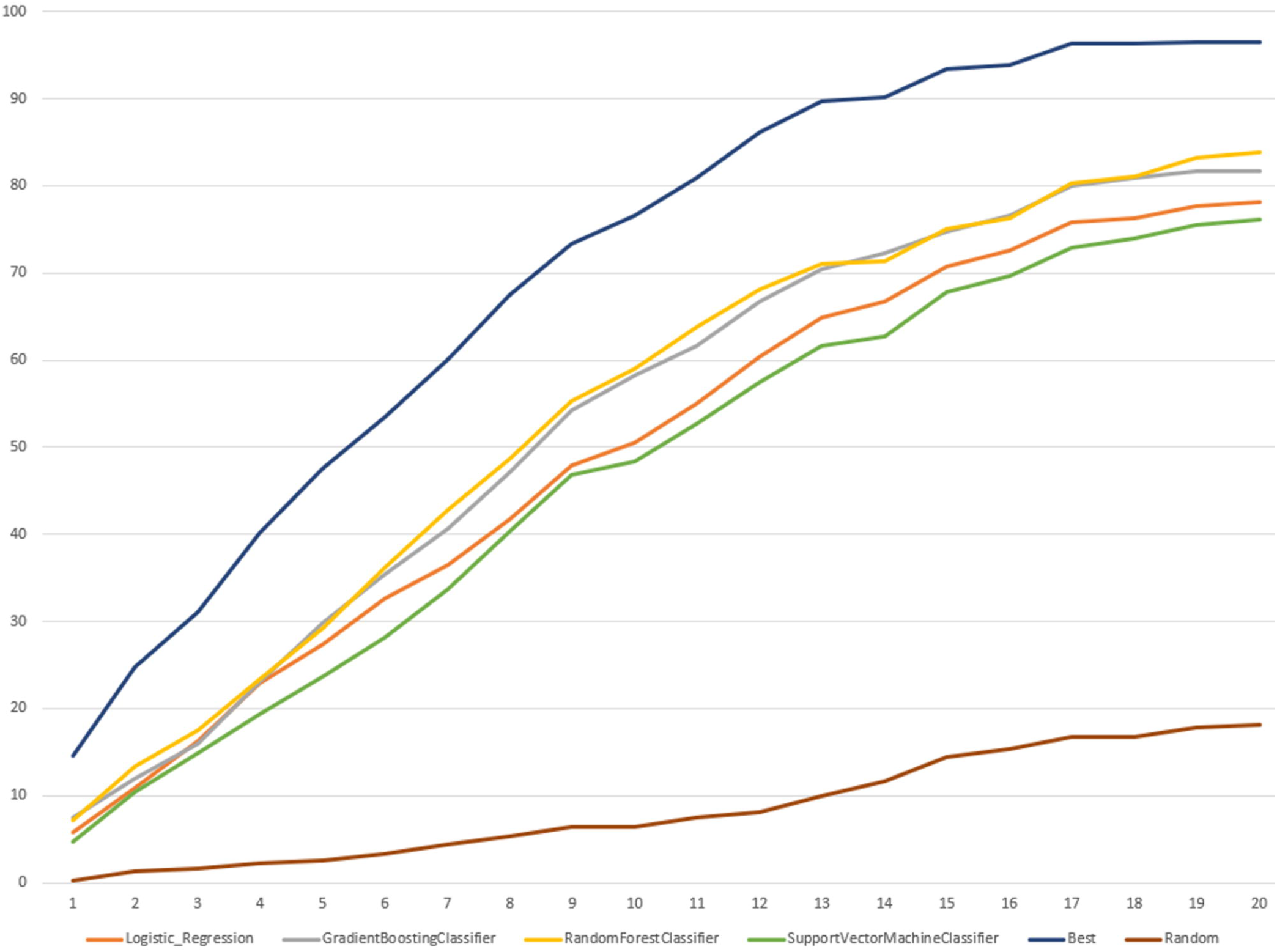
The cumulative ranking score in percentage for the recovered known sites of cleavage for all proteases in range of ranking positions from 0 to 20%.

All classifiers were able to recover all known cleavage sites better than random but at lower percentage of ranking positions than the best. LR models were the best considering they recovered all known SoCs at 99% of the ranking positions, while the best recovered known sites of cleavage at 31%. RFC and GBC recovered all SoCs from external dataset for all proteases at 91% of the ranking positions, accordingly. Moreover, we compared percentage of recovered known SoCs at the percentage ranking positions when the best ranking collected 100% of known sites of cleavage for each protease family selected for this study, results are presented in S9 Table. For all protease families except of cysteine the highest percentage of correctly predicted cleavage sites was reached by RCF models and the lowest by SVC. S10 Table summarizes the normalized ranking positions reached when 100% of known cleavage sites were recovered for all classifiers for all selected protease families.

Finally, we compared the predictive performance by ranking on external dataset between LR and RFC classifiers and other tools such as SitePrediction and PROSPERous. For following proteases predictive performance was compared with PROSPERous and SitePrediction: caspase-1, caspase-3, caspase-6, caspase-7, cathepsin E, cathepsin D, cathepsin L, MMP2, MMP3, MMP8, MMP9. Also, PROSPERous contained models trained for serine proteases: granzyme A, granzyme B, granzyme B (rodent-type), thrombin. Our LR and RFC models as well as PROSPERous correctly predicted 376 known cleavage sites, while 306 known cleavage sites were correctly predicted by both PROSPERous and SitePrediction. Fig 8 and Fig 9 summarizes the cumulative percentage score for the recovered known sites of cleavage between our all models compared to PROSPERous and SitePrediction performance on external dataset.

**Figure 8.**
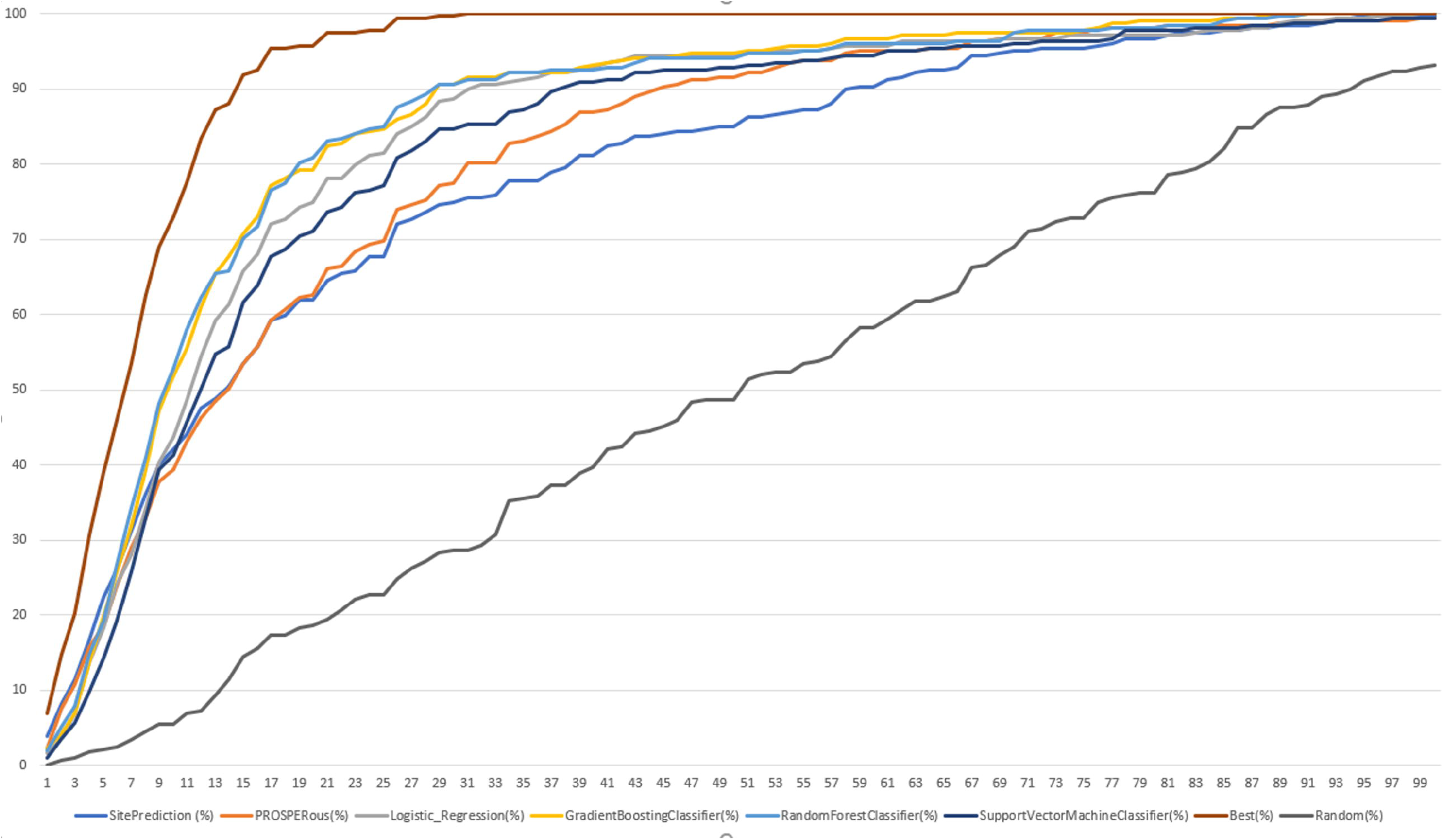
The cumulative ranking score in percentage for the recovered known sites of cleavage for selected protease families compared with PROSPERous and SitePrediction.

**Figure 9.**
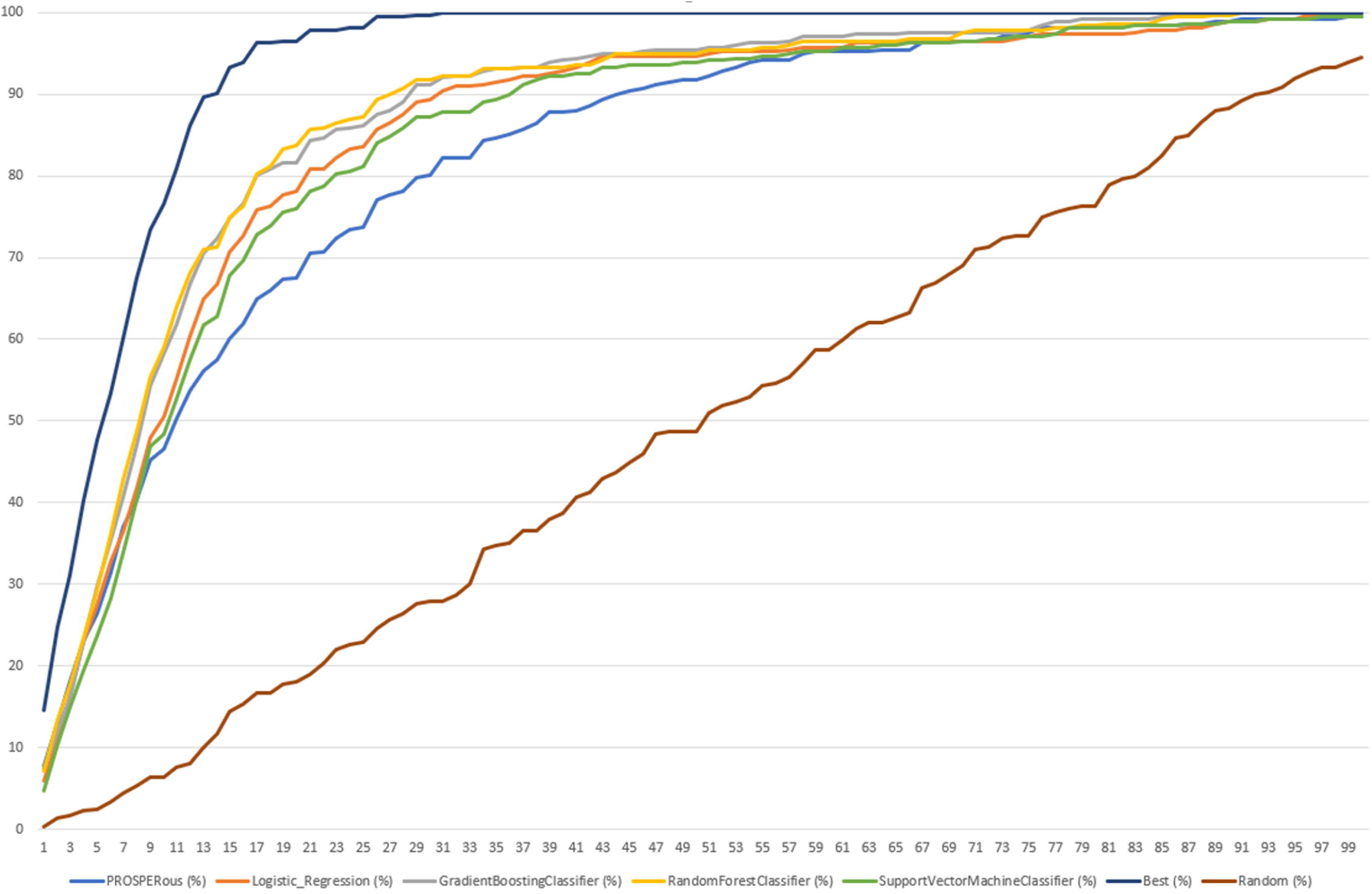
The cumulative ranking score in percentage for the recovered known sites of cleavage for selected protease families compared with PROSPERous.

Our models were able to collect the same amount of known cleavage sites as PROSPERous and SitePrediction. SitePrediction is based on statistical scoring methods, while PROSPERous is based on SVM learning and LR approach. In the dataset of proteases compared to SitePrediction and PROSPERous LR and RFC models recovered higher percentage of known SoCs at the ranking percentage position equal to 50% but reaching 88% the percentage of recovered known sites of cleavage was the same for our models, PROSPERous and SitePrediction. In the dataset of proteases compared only to PROSPERous RFC model performed worse than PROSPERous and recovered less percentage of known cleavage sites. On the other hand, LR model recovered higher percentage of known SoCs at the ranking percentage position equals to 50% but reaching about 85% of ranking position percentage of recovered known sites of cleavage was the same for LR and PROSPERous. While we performed detailed analysis of ranking position percentage between 0 and 40% LR outperformed RFC and PROSPERous, while last ones performed at the same level. We can suppose that LR model works better for the peptides of the smaller size.

## Conclusions

Predicting possible sites of cleavage for individual proteases is an important task to be completed during drug-design process of peptide therapeutics to improve their stability and availably as a promising drug. In this study we presented a new approach in WebMetabase that helps to predict cleavage site for the specific peptide family or for specific experimental condition (i.e. individual protease). One of the main advantages of this approach is that it generates a searchable database for the information coming from LC-MS based experimental data or from external sources such as MEROPS database. In this database each amino acid is described as a vector of physicochemical properties, Volsurf molecular descriptors. Thus, the sequence pattern around the potential cleavage was represented as combination of Volsurf descriptors. The proposed methodology can be applied in the case of non-natural amino acid. Comparing to MEROPS this database type can be enriched with new experimental or external data. This way to store the data can be utilized to perform frequency analysis to discover the most frequent scissile bonds within the generated database. The FA results can be used to derive a cleavage site appearance rules based on molecular properties of the cleavage sites. To demonstrate this, we trained several models using Logistic Regression (LR), Support Vector Machine (SVM) and Ensemble Trees (ET) classifier learning approaches for eighteen proteases from four protease families: serine, cysteine, aspartic and matrix metalloproteases. In the training dataset each sequence pattern around the potential cleavage site and actual site of cleavage was represented as a combination of Volsurf descriptors that characterized physicochemical properties of the SBs in the sequence. We compared predictive performance of the models trained with different learning approaches applying 5-fold cross validation test and external dataset validation test. Moreover, we examined the influence of the local window sequence size around the site of cleavage by comparing the models trained for P1-P1’ and P4-P4’ range. We revealed that LR and RFC models trained using window P4-P4’ outperformed other learning methods and the models trained using P1-P1’ window. We noted that training dataset size influenced on the predictive performance of the models analyzing data for caspases. Finally, we compared the predictive performance of LR and RFC models with other approaches such as PROSPERous and SitePrediction tools. LR model recovered higher percentage of the known cleavage site in the first 30% of the ranking positions comparing to the other tools. It can be explained by the fact that it performs better prediction on smaller peptides. Moreover, since the system can be linked to the software assisted metabolite structure elucidation based on MS data, the database is automatically enriched with the new experiments. Nevertheless, models can be re-trained with updated dataset and derived rules can be refined to tune the system for the experimental conditions and/or peptide families of interest. This knowledge can be applied during the design-make-test drug discovery cycle.

## Supporting information

## Acknowledgments

This work has been partially supported by Doctorats Industrials, AGAUR, Generalitat de Catalunya. We would like to thank Xavier Pascual from Lead Molecular Design for the programming support and Dr. Manuel Pastor from Barcelona Biomedical Research Park, Universidad Pompeu Fabra for his contributions and help in statistical analysis discussions.

## Supporting information

**S1 Table. Summary on the extracted information from MEROPS database.**

**S2 Table. Summary on the model’s number for each protease for cleavage window P1-P1’ and P4-P4’.**

**S3 Table. Summary on count of individual SoC2 and SoC8 calculated by frequency analysis.**

**S4 Table. Results for the predictive performance evaluation for LR and SVC classifiers trained using local windows P1-P1’ and P4-P4’ for caspases all evaluated parameters for 5fold CV.**

**S5 Table. Results for the predictive performance evaluation for LR and SVC classifiers trained using local windows P1-P1’ and P4-P4’ for caspases all evaluated parameters for external dataset.**

**S6 Table. Results for the predictive performance evaluation for LR and SVC classifiers validated on external dataset for all selected proteases and all evaluated parameters trained using local window P1-P1’.**

**S7 Table. Results for the predictive performance evaluation for LR and SVC classifiers validated on external dataset for all selected proteases and all evaluated parameters trained using local window P4-P4’.**

**S8 Table. Performance metrics calculated during validation on external dataset for all investigated classifier models trained using window P4-P4’ for all selected proteases.**

**S9 Table. The percentage of recovered known SoCs at the percentage ranking positions when the best ranking collected 100% of known sites of cleavage for each protease family.**

**S10 Table. The normalized ranking positions reached when 100% of known cleavage sites were recovered for all classifiers for all selected protease families.**

**S1 Figure. Substrate cleavage sites for P4-P4’ local window for cathepsins: a) cathepsin D; b) cathepsin E; c) cathepsin L. The cumulative amino acid occurrences in P4–P4’ were calculated and displayed in the form of a two-dimensional heat map. The scissile peptide bond was between sites P1 and P1’.**

**S2 Figure. Substrate cleavage sites for P4–P4’ position for granzymes. a) granzyme A; b) granzyme B; c) granzyme B (rodent-type); d) granzyme M. The cumulative amino acid occurrences in P4–P4’ were calculated and displayed in the form of a two-dimensional heat map. The scissile peptide bond was between sites P1 and P1’.**

**S3 Figure. Substrate cleavage sites for P4–P4’ position for matrix metalloproteases (MMP). a) MMP2; b) MMP3; c) MMP8; d) MMP9. The cumulative amino acid occurrences in P4–P4’ were calculated and displayed in the form of a two-dimensional heat map. The scissile peptide bond was between sites P1 and P1’.**

**S1 File. Information about cleaved and noncleaved motifs from for all selected proteases in the training datasets and external datasets described by Volsurf descriptors.**

**S2 File. Python script for model generation and ReadMe.txt with script usage instruction.**

**S3 File. Python script for the prediction with generated models and ReadMe.txt with script usage instruction.**

