## Supplementary figures and images for "Software-aided workflow for predicting protease-specific cleavage sites using physicochemical properties of the natural and unnatural amino acids in peptide-based drug discovery: Peptide cleavage sites prediction workflow"

### Supplementary file 1

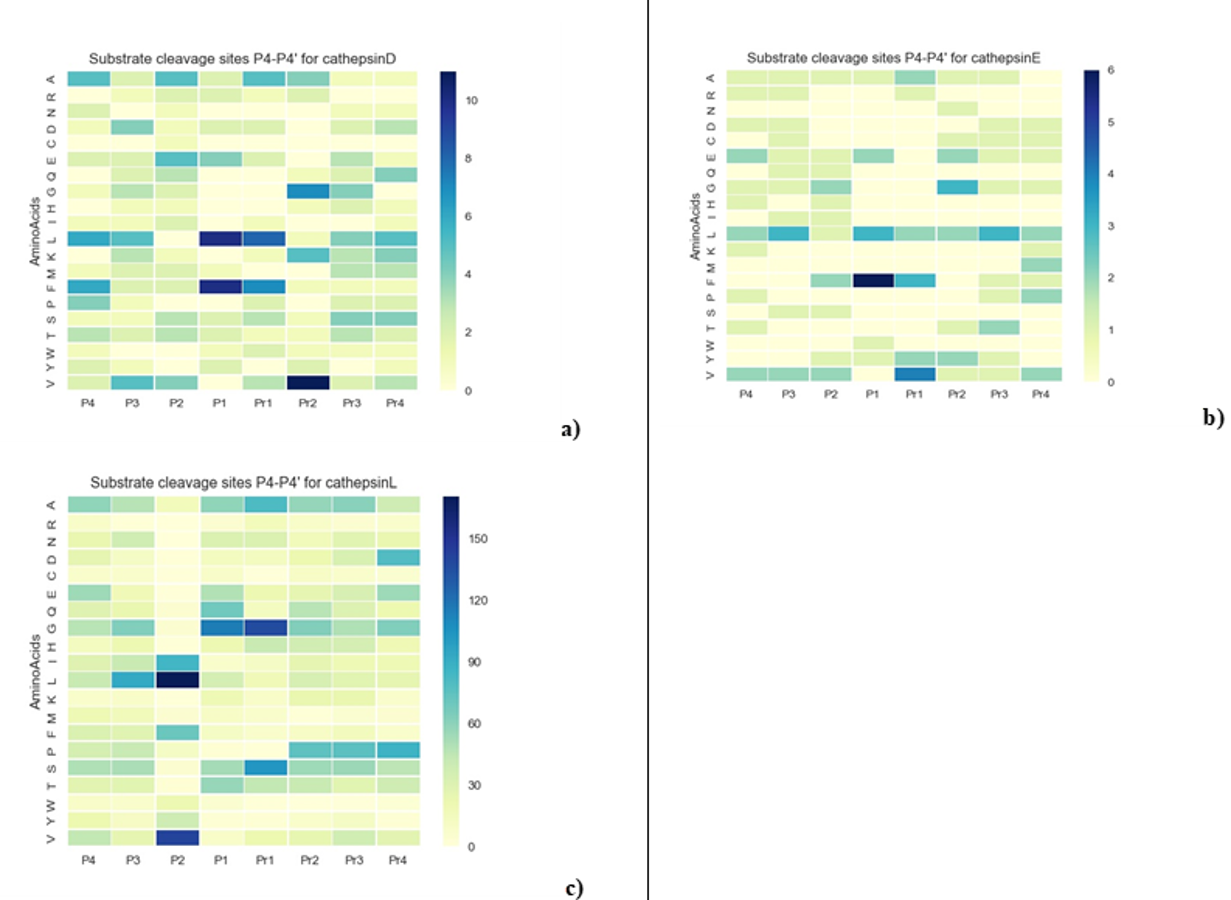

### Supplementary file 3

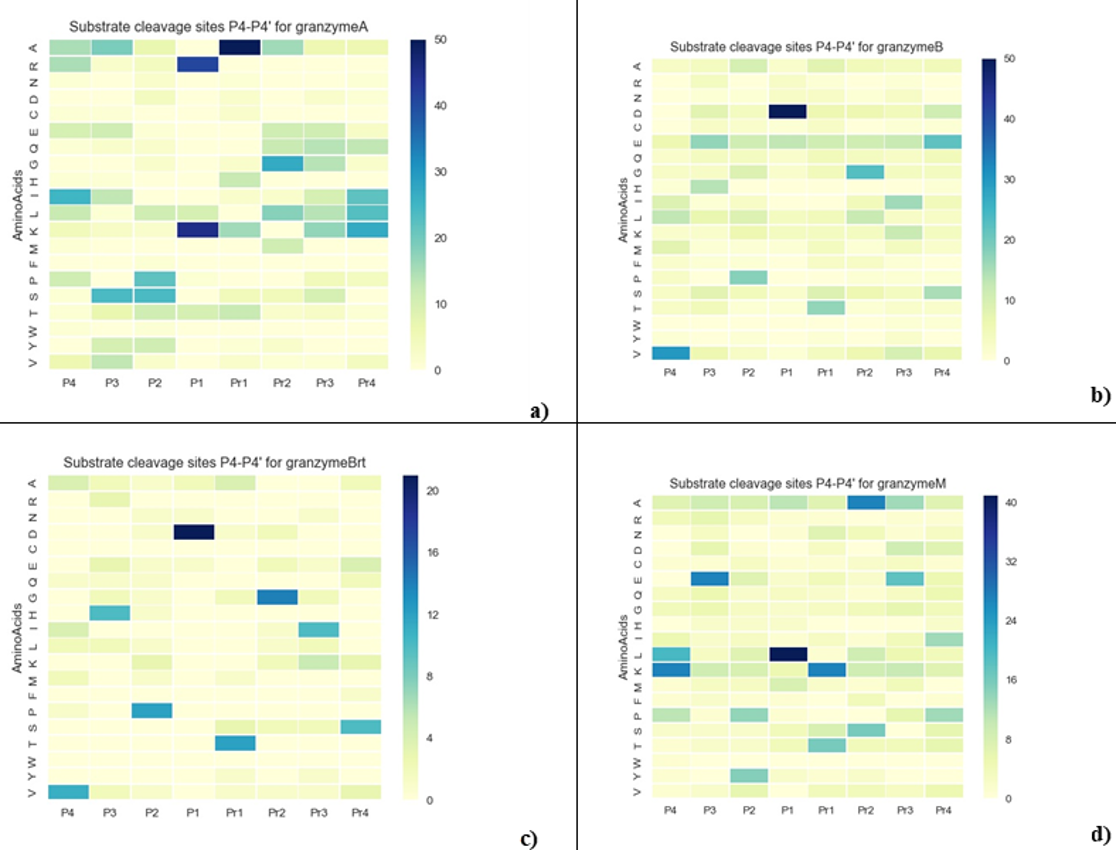

### Supplementary file 5

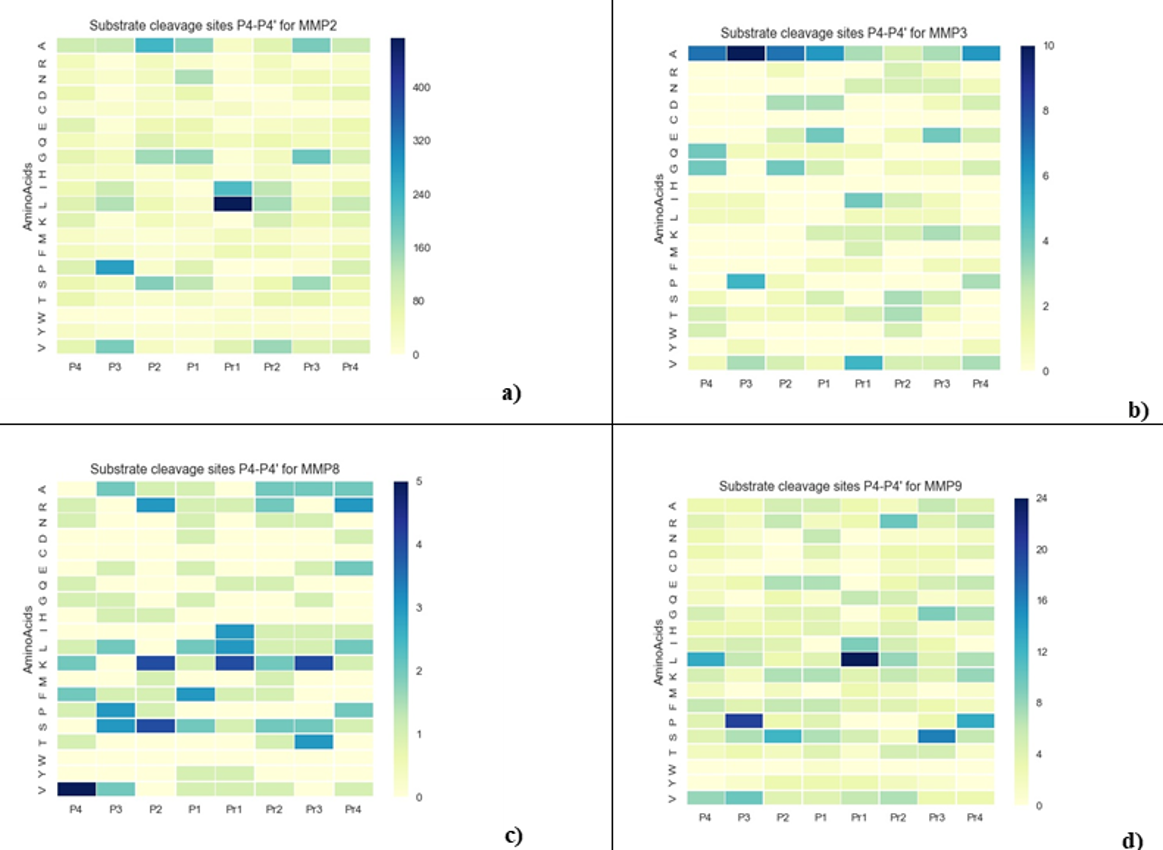
