## Supplementary material for "Software-aided workflow for predicting protease-specific cleavage sites using physicochemical properties of the natural and unnatural amino acids in peptide-based drug discovery: Peptide cleavage sites prediction workflow"

### Supporting Table 1. Summary on the extracted information from MEROPS database

| Protease family | Protease | Substrates | Cleavage events | SoC2 P1-P1’ | SoC8 P4-P4’ | Substrates in external dataset |
|---|---|---|---|---|---|---|
| Serine proteases | Granzyme B (rodent-type) | 31 | 32 | 11 | 15 | 7 |
| | Trypsin 1 | 327 | 1104 | 43 | 847 | 31 |
| | Granzyme M | 72 | 164 | 47 | 70 | 7 |
| | Granzyme A | 66 | 135 | 18 | 40 | 14 |
| | Granzyme B | 65 | 121 | 46 | 73 | 17 |
| | Thrombin | 53 | 59 | 14 | 39 | 12 |
| Matrix metallopeptidases | Matrix Metallopeptidase-2 | 1359 | 1528 | 199 | 1057 | 53 |
| | Matrix Metallopeptidase-3 | 11 | 29 | 21 | 23 | 4 |
| | Matrix Metallopeptidase-8 | 11 | 25 | 16 | 16 | 3 |
| | Matrix Metallopeptidase-9 | 20 | 185 | 58 | 74 | 2 |
| Aspartic proteases | Cathepsin D | 16 | 53 | 37 | 38 | 5 |
| | Cathepsin E | 8 | 20 | 11 | 14 | 3 |
| Cysteine Aspartate proteases | Caspase-1 | 10 | 10 | 6 | 8 | 2 |
| | Caspase-2 | 216 | 221 | 17 | 136 | 44 |
| | Caspase-3 | 52 | 72 | 13 | 42 | 10 |
| | Caspase-6 | 870 | 885 | 25 | 782 | 10 |
| | Caspase-7 | 23 | 29 | 6 | 13 | 6 |
| Cysteine proteases | Cathepsin L | 895 | 964 | 209 | 587 | 186 |
