## Supplementary material for "Software-aided workflow for predicting protease-specific cleavage sites using physicochemical properties of the natural and unnatural amino acids in peptide-based drug discovery: Peptide cleavage sites prediction workflow"

**Supporting Table 2. Summary on the model's number for each protease for cleavage window P1-P1' and P4-P4'**

| Protease | Models number for SoC2 | Models number for SoC8 |
|---|---|---|
| **Granzyme B (rodent-type)** | 32 | 126 |
| **Trypsin 1** | 13 | 38 |
| **Granzyme M** | 8 | 79 |
| **Granzyme A** | 21 | 94 |
| **Granzyme B** | 8 | 102 |
| **Thrombin** | 24 | 21 |
| **Matrix Metallopeptidase-2** | 2 | 14 |
| **Matrix Metallopeptidase-3** | 11 | 26 |
| **Matrix Metallopeptidase-8** | 15 | 38 |
| **Matrix Metallopeptidase-9** | 4 | 16 |
| **Cathepsin D** | 7 | 28 |
| **Cathepsin E** | 7 | 5 |
| **Caspase-1** | 57 | 148 |
| **Caspase-2** | 27 | 25 |
| **Caspase-3** | 31 | 54 |
| **Caspase-6** | 25 | 23 |
| **Caspase-7** | 33 | 27 |
| **Cathepsin L** | 2 | 14 |
