## Supplementary material for "Software-aided workflow for predicting protease-specific cleavage sites using physicochemical properties of the natural and unnatural amino acids in peptide-based drug discovery: Peptide cleavage sites prediction workflow"

**Supporting Table 3. Summary on count of individual SoC2 and SoC8 calculated by frequency analysis**

| Protease | Datasets for window P1-P1’ | | | | Datasets for window P4-P4’ | | | |
|---|---|---|---|---|---|---|---|---|
| | Train dataset size | | External dataset size | | Train dataset size | | External dataset size | |
| | Negative SoCs | Positive SoCs | Negative SoCs | Positive SoCs | Negative SoCs | Positive SoCs | Negative SoCs | Positive SoCs |
| **Granzyme B (rodent-type)** | 371 | 11 | 270 | 4 | 1923 | 15 | 920 | 7 |
| **Trypsin 1** | 633 | 43 | 434 | 33 | 34217 | 847 | 4031 | 113 |
| **Granzyme M** | 458 | 47 | 313 | 11 | 5676 | 70 | 956 | 18 |
| **Granzyme A** | 420 | 18 | 342 | 10 | 3856 | 40 | 1910 | 27 |
| **Granzyme B** | 471 | 46 | 396 | 15 | 7558 | 73 | 2323 | 24 |
| **Thrombin** | 363 | 14 | 146 | 8 | 915 | 39 | 154 | 11 |
| **Matrix Metallopeptidase-2** | 718 | 199 | 424 | 48 | 17017 | 1057 | 1059 | 54 |
| **Matrix Metallopeptidase-3** | 278 | 21 | 242 | 6 | 651 | 23 | 362 | 6 |
| **Matrix Metallopeptidase-8** | 288 | 16 | 85 | 5 | 636 | 16 | 79 | 2 |
| **Matrix Metallopeptidase-9** | 376 | 58 | 211 | 19 | 1365 | 74 | 400 | 21 |
| **Cathepsin D** | 348 | 37 | 213 | 11 | 1129 | 38 | 346 | 11 |
| **Cathepsin E** | 102 | 11 | 192 | 6 | 94 | 14 | 264 | 6 |
| **Caspase-1** | 356 | 6 | 188 | 2 | 1200 | 8 | 312 | 2 |
| **Caspase-2** | 491 | 17 | 320 | 7 | 3610 | 136 | 961 | 36 |
| **Caspase-3** | 435 | 13 | 356 | 7 | 2361 | 42 | 1421 | 14 |
| **Caspase-6** | 668 | 25 | 259 | 6 | 19229 | 782 | 574 | 11 |
| **Caspase-7** | 212 | 6 | 234 | 2 | 382 | 13 | 417 | 5 |
| **Cathepsin L** | 765 | 209 | 576 | 118 | 9479 | 587 | 2726 | 174 |
