## Supplementary material for "Software-aided workflow for predicting protease-specific cleavage sites using physicochemical properties of the natural and unnatural amino acids in peptide-based drug discovery: Peptide cleavage sites prediction workflow"

**S4 Table. The predictive performance evaluation for Logistic Regression and Support Vector Machine Classifiers based on results of the 5-fold Cross validation for caspases and cathepsin L.**

| | | Cleavage site size P1-P1' | | | | | | Cleavage site size P4-P4' | | | | | |
|---|---|---|---|---|---|---|---|---|---|---|---|---|---|
| | | caspase1 | caspase2 | caspase3 | caspase6 | caspase7 | cathepsinL | caspase1 | caspase2 | caspase3 | caspase6 | caspase7 | cathepsinL |
| Accuracy | LR | 0.77 | 0.89 | 0.90 | 0.94 | 0.77 | 0.83 | 0.73 | 0.93 | 0.88 | 0.96 | 0.87 | 0.84 |
| | SVC | 0.89 | 0.90 | 0.90 | 0.93 | 0.83 | 0.81 | 0.67 | 0.86 | 0.79 | 0.90 | 0.68 | 0.77 |
| AUC ROC | LR | 0.77 | 0.87 | 0.90 | 0.92 | 0.77 | 0.81 | 0.73 | 0.93 | 0.88 | 0.96 | 0.87 | 0.84 |
| | SVC | 0.89 | 0.89 | 0.91 | 0.92 | 0.83 | 0.76 | 0.67 | 0.86 | 0.79 | 0.90 | 0.68 | 0.77 |
| MCC | LR | 0.58 | 0.78 | 0.82 | 0.86 | 0.58 | 0.62 | 0.50 | 0.87 | 0.77 | 0.92 | 0.76 | 0.69 |
| | SVC | 0.80 | 0.81 | 0.82 | 0.86 | 0.69 | 0.54 | 0.37 | 0.74 | 0.63 | 0.80 | 0.36 | 0.54 |
| AUC PRC | LR | 0.73 | 0.87 | 0.89 | 0.93 | 0.76 | 0.84 | 0.72 | 0.90 | 0.83 | 0.94 | 0.84 | 0.79 |
| | SVC | 0.87 | 0.88 | 0.90 | 0.94 | 0.83 | 0.83 | 0.67 | 0.85 | 0.78 | 0.88 | 0.68 | 0.72 |
| Sensitivity | LR | 0.96 | 0.95 | 0.97 | 0.98 | 0.92 | 0.88 | 0.80 | 0.96 | 0.93 | 0.97 | 0.95 | 0.82 |
| | SVC | 0.92 | 0.90 | 0.91 | 0.94 | 0.83 | 0.87 | 0.55 | 0.76 | 0.62 | 0.84 | 0.37 | 0.68 |
| Specificity | LR | 0.59 | 0.79 | 0.82 | 0.86 | 0.62 | 0.73 | 0.66 | 0.90 | 0.83 | 0.95 | 0.79 | 0.87 |
| | SVC | 0.85 | 0.87 | 0.90 | 0.91 | 0.83 | 0.65 | 0.78 | 0.96 | 0.97 | 0.96 | 0.98 | 0.86 |
