## Supplementary material for "Software-aided workflow for predicting protease-specific cleavage sites using physicochemical properties of the natural and unnatural amino acids in peptide-based drug discovery: Peptide cleavage sites prediction workflow"

**S5 Table. The predictive performance evaluation for Logistic Regression and Support Vector Machine Classifiers based on results of the external validation for caspases and cathepsin L.**

| Learning algorithm | Local window size | Performance metrics | caspase1 | caspase2 | caspase3 | caspase6 | caspase7 | cathepsinL |
|---|---|---|---|---|---|---|---|---|
| **LR** | **P1/P1'** | **Accuracy** | 0.68 | 0.83 | 0.80 | 0.82 | 0.70 | 0.33 |
| | | **AUC PRC** | 0.14 | 0.49 | 0.23 | 0.40 | 0.25 | 0.14 |
| | | **AUC ROC** | 0.03 | 0.31 | 0.08 | 0.22 | 0.10 | 0.09 |
| | | **MCC** | 0.84 | 0.91 | 0.87 | 0.91 | 0.83 | 0.62 |
| | | **Sensitivity** | 1.00 | 1.00 | 0.94 | 1.00 | 0.97 | 0.95 |
| | | **Specificity** | 0.68 | 0.82 | 0.79 | 0.81 | 0.69 | 0.29 |
| | **P4/P4'** | **Accuracy** | 0.69 | 0.76 | 0.84 | 0.75 | 0.64 | 0.72 |
| | | **AUC PRC** | 0.10 | 0.53 | 0.22 | 0.46 | 0.22 | 0.41 |
| | | **AUC ROC** | 0.02 | 0.43 | 0.09 | 0.36 | 0.10 | 0.33 |
| | | **MCC** | 0.77 | 0.77 | 0.84 | 0.74 | 0.68 | 0.73 |
| | | **Sensitivity** | 0.86 | 0.79 | 0.85 | 0.74 | 0.73 | 0.73 |
| | | **Specificity** | 0.69 | 0.75 | 0.83 | 0.75 | 0.64 | 0.73 |
| **SVC** | **P1/P1'** | **Accuracy** | 0.83 | 0.85 | 0.97 | 0.83 | 0.85 | 0.33 |
| | | **AUC PRC** | 0.21 | 0.50 | 0.85 | 0.40 | 0.38 | 0.13 |
| | | **AUC ROC** | 0.05 | 0.33 | 0.27 | 0.22 | 0.20 | 0.09 |
| | | **MCC** | 0.92 | 0.92 | 0.10 | 0.91 | 0.90 | 0.62 |
| | | **Sensitivity** | 1.00 | 1.00 | 0.91 | 1.00 | 0.96 | 0.94 |
| | | **Specificity** | 0.83 | 0.84 | 0.84 | 0.82 | 0.84 | 0.29 |
| | **P4/P4'** | **Accuracy** | 0.74 | 0.77 | 0.88 | 0.76 | 0.72 | 0.73 |
| | | **AUC PRC** | 0.12 | 0.55 | 0.26 | 0.50 | 0.29 | 0.37 |
| | | **AUC ROC** | 0.02 | 0.46 | 0.11 | 0.39 | 0.17 | 0.32 |
| | | **MCC** | 0.82 | 0.77 | 0.86 | 0.74 | 0.71 | 0.69 |
| | | **Sensitivity** | 0.90 | 0.78 | 0.83 | 0.72 | 0.70 | 0.63 |
| | | **Specificity** | 0.74 | 0.77 | 0.88 | 0.76 | 0.72 | 0.74 |
