## Supplementary material for "Software-aided workflow for predicting protease-specific cleavage sites using physicochemical properties of the natural and unnatural amino acids in peptide-based drug discovery: Peptide cleavage sites prediction workflow"

**S8 Table.The performance validation results for all classifiers on external dataset for P4/P4' models.**

| Performance metrics | Learning algorithm | Accuracy | AUC PRC | AUC ROC | MCC | Sensitivity | Specificity |
|---|---|---|---|---|---|---|---|
| **Caspase1** | **LR** | 0.69 | 0.10 | 0.02 | 0.77 | 0.86 | 0.69 |
| | **SVC** | 0.74 | 0.12 | 0.02 | 0.82 | 0.90 | 0.74 |
| | **RFC** | 0.85 | 0.18 | 0.05 | 0.88 | 0.91 | 0.85 |
| | **GBC** | 0.85 | 0.21 | 0.05 | 0.91 | 0.97 | 0.85 |
| **Caspase2** | **LR** | 0.76 | 0.53 | 0.43 | 0.77 | 0.79 | 0.75 |
| | **SVC** | 0.77 | 0.55 | 0.46 | 0.77 | 0.78 | 0.77 |
| | **RFC** | 0.77 | 0.58 | 0.48 | 0.79 | 0.81 | 0.77 |
| | **GBC** | 0.77 | 0.57 | 0.48 | 0.79 | 0.81 | 0.77 |
| **Caspase3** | **LR** | 0.84 | 0.22 | 0.09 | 0.84 | 0.85 | 0.83 |
| | **SVC** | 0.88 | 0.26 | 0.11 | 0.86 | 0.83 | 0.88 |
| | **RFC** | 0.94 | 0.36 | 0.19 | 0.89 | 0.84 | 0.94 |
| | **GBC** | 0.94 | 0.36 | 0.19 | 0.89 | 0.84 | 0.94 |
| **Caspase6** | **LR** | 0.75 | 0.46 | 0.36 | 0.74 | 0.74 | 0.75 |
| | **SVC** | 0.76 | 0.50 | 0.39 | 0.74 | 0.72 | 0.76 |
| | **RFC** | 0.76 | 0.46 | 0.32 | 0.74 | 0.72 | 0.76 |
| | **GBC** | 0.76 | 0.47 | 0.32 | 0.76 | 0.76 | 0.76 |
| **Caspase7** | **LR** | 0.64 | 0.22 | 0.10 | 0.68 | 0.73 | 0.64 |
| | **SVC** | 0.72 | 0.29 | 0.17 | 0.71 | 0.70 | 0.72 |
| | **RFC** | 0.77 | 0.38 | 0.24 | 0.77 | 0.77 | 0.77 |
| | **GBC** | 0.73 | 0.35 | 0.21 | 0.76 | 0.79 | 0.73 |
| **CathepsinD** | **LR** | 0.64 | 0.08 | 0.06 | 0.62 | 0.60 | 0.64 |
| | **SVC** | 0.64 | 0.07 | 0.06 | 0.61 | 0.57 | 0.64 |
| | **RFC** | 0.70 | 0.16 | 0.08 | 0.70 | 0.71 | 0.70 |
| | **GBC** | 0.66 | 0.14 | 0.07 | 0.69 | 0.73 | 0.66 |
| **CathepsinE** | **LR** | 0.58 | 0.09 | 0.05 | 0.67 | 0.76 | 0.57 |
| | **SVC** | 0.57 | 0.09 | 0.05 | 0.67 | 0.78 | 0.56 |
| | **RFC** | 0.71 | 0.15 | 0.06 | 0.74 | 0.78 | 0.71 |
| | **GBC** | 0.63 | 0.14 | 0.06 | 0.73 | 0.83 | 0.62 |
| **CathepsinL** | **LR** | 0.72 | 0.41 | 0.33 | 0.73 | 0.73 | 0.73 |
| | **SVC** | 0.73 | 0.37 | 0.32 | 0.69 | 0.63 | 0.74 |
| | **RFC** | 0.75 | 0.45 | 0.37 | 0.74 | 0.73 | 0.75 |
| | **GBC** | 0.75 | 0.47 | 0.38 | 0.76 | 0.78 | 0.74 |
| **GranzymeA** | **LR** | 0.73 | 0.17 | 0.05 | 0.83 | 0.93 | 0.73 |
| | **SVC** | 0.77 | 0.19 | 0.06 | 0.85 | 0.93 | 0.77 |
| | **RFC** | 0.85 | 0.26 | 0.09 | 0.90 | 0.96 | 0.84 |
| | **GBC** | 0.80 | 0.22 | 0.07 | 0.87 | 0.95 | 0.80 |
| **GranzymeB** | **LR** | 0.69 | 0.12 | 0.04 | 0.77 | 0.85 | 0.69 |
| | **SVC** | 0.73 | 0.13 | 0.04 | 0.79 | 0.84 | 0.73 |
| | **RFC** | 0.80 | 0.16 | 0.05 | 0.83 | 0.86 | 0.80 |
| | **GBC** | 0.74 | 0.13 | 0.04 | 0.81 | 0.87 | 0.74 |
| **GranzymeBrt** | **LR** | 0.65 | 0.12 | 0.02 | 0.83 | 1.00 | 0.65 |
| | **SVC** | 0.71 | 0.14 | 0.03 | 0.85 | 1.00 | 0.71 |
| | **RFC** | 0.84 | 0.21 | 0.06 | 0.92 | 1.00 | 0.84 |
| | **GBC** | 0.73 | 0.15 | 0.03 | 0.86 | 1.00 | 0.72 |

| S8 Table.The performance validation results for all classifiers on external dataset for P4/P4' models. | | | | | | | |
|---|---|---|---|---|---|---|---|
| **Performance metrics** | **Learning algorithm** | **Accuracy** | **AUC PRC** | **AUC ROC** | **MCC** | **Sensitivity** | **Specificity** |
| **GranzymeM** | **LR** | 0.66 | 0.14 | 0.04 | 0.76 | 0.86 | 0.66 |
| | **SVC** | 0.68 | 0.15 | 0.05 | 0.77 | 0.85 | 0.68 |
| | **RFC** | 0.78 | 0.19 | 0.06 | 0.80 | 0.82 | 0.78 |
| | **GBC** | 0.73 | 0.19 | 0.06 | 0.81 | 0.90 | 0.73 |
| **MMP2** | **LR** | 0.78 | 0.49 | 0.38 | 0.79 | 0.82 | 0.77 |
| | **SVC** | 0.80 | 0.43 | 0.37 | 0.73 | 0.65 | 0.81 |
| | **RFC** | 0.78 | 0.47 | 0.37 | 0.78 | 0.78 | 0.77 |
| | **GBC** | 0.77 | 0.51 | 0.38 | 0.80 | 0.84 | 0.78 |
| **MMP3** | **LR** | 0.59 | 0.10 | 0.12 | 0.62 | 0.66 | 0.57 |
| | **SVC** | 0.72 | 0.20 | 0.19 | 0.69 | 0.66 | 0.72 |
| | **RFC** | 0.76 | 0.17 | 0.16 | 0.68 | 0.57 | 0.78 |
| | **GBC** | 0.69 | 0.12 | 0.13 | 0.64 | 0.57 | 0.71 |
| **MMP8** | **LR** | 0.41 | 0.16 | 0.15 | 0.49 | 0.58 | 0.39 |
| | **SVC** | 0.44 | 0.19 | 0.18 | 0.50 | 0.57 | 0.43 |
| | **RFC** | 0.42 | 0.12 | 0.15 | 0.45 | 0.49 | 0.42 |
| | **GBC** | 0.38 | 0.01 | 0.12 | 0.39 | 0.38 | 0.40 |
| **MMP9** | **LR** | 0.59 | 0.09 | 0.11 | 0.60 | 0.62 | 0.59 |
| | **SVC** | 0.59 | 0.09 | 0.10 | 0.60 | 0.61 | 0.59 |
| | **RFC** | 0.70 | 0.16 | 0.13 | 0.68 | 0.66 | 0.71 |
| | **GBC** | 0.64 | 0.16 | 0.12 | 0.69 | 0.74 | 0.64 |
| **Thrombin** | **LR** | 0.85 | 0.69 | 0.58 | 0.88 | 0.92 | 0.84 |
| | **SVC** | 0.85 | 0.69 | 0.59 | 0.88 | 0.92 | 0.84 |
| | **RFC** | 0.88 | 0.77 | 0.70 | 0.89 | 0.91 | 0.87 |
| | **GBC** | 0.84 | 0.67 | 0.58 | 0.86 | 0.90 | 0.83 |
| **Trypsin1** | **LR** | 0.88 | 0.40 | 0.21 | 0.88 | 0.89 | 0.88 |
| | **SVC** | 0.89 | 0.36 | 0.20 | 0.79 | 0.69 | 0.90 |
| | **RFC** | 0.87 | 0.41 | 0.22 | 0.90 | 0.92 | 0.87 |
| | **GBC** | 0.87 | 0.40 | 0.22 | 0.90 | 0.92 | 0.87 |
