## Supplementary material for "Software-aided workflow for predicting protease-specific cleavage sites using physicochemical properties of the natural and unnatural amino acids in peptide-based drug discovery: Peptide cleavage sites prediction workflow"

**Supporting Table 9. The percentage of recovered known cleavage sites at percentage of ranking position reached when best recovered 100% of all known sites of cleavage for all selected proteases. Highest recovered percentage marked in bold.**

| | LR (%) | SVC (%) | RCF (%) | GBC (%) | Best (%) | Random (%) |
|---|---|---|---|---|---|---|
| **Serine protease** | 93 | 91 | **96** | **96** | 100.0 | 13 |
| **Cysteine protease** | 94 | 89 | 94 | **95** | 100.0 | 29 |
| **Aspartic protease** | 29 | 29 | **47** | 35 | 100.0 | 0 |
| **Matrix metalloproteases** | **86** | 82 | **86** | 83 | 100.0 | 25 |
