## Supplementary material for "Software-aided workflow for predicting protease-specific cleavage sites using physicochemical properties of the natural and unnatural amino acids in peptide-based drug discovery: Peptide cleavage sites prediction workflow"

**Supporting Table 10. The normalized ranking percentage position reached by logistic regression and random forest models when 100% of known SoC was recovered for all selected proteases. Lowest ranking percentage position marked in bold.**

| | **LR (%)** | **SVC (%)** | **RCF (%)** | **GBC (%)** | **Best (%)** | **Random (%)** |
|---|---|---|---|---|---|---|
| **Serine protease** | 95 | **73** | 86 | 88 | 12 | 100 |
| **Cysteine protease** | 95 | 100 | 88 | **84** | 30 | 100 |
| **Aspartic protease** | 98 | 96 | **84** | 90 | 7 | 87 |
| **Matrix metalloproteases** | 89 | 100 | 90 | **87** | 25 | 100 |
